# Myeloid Drp1 Deficiency Limits Revascularization in Ischemic Muscles via Inflammatory Macrophage Polarization and Metabolic Reprograming

**DOI:** 10.1101/2023.11.04.565656

**Authors:** Shikha Yadav, Vijay Ganta, Varadarajan Sudhahar, Dipankar Ash, Sheela Nagarkoti, Archita Das, Margorzata McMenamin, Stephanie Kelley, Tohru Fukai, Masuko Ushio-Fukai

**Affiliations:** Vascular Biology Center; Department of Medicine (Cardiology); Department of Pharmacology and Toxicology, Medical College of Georgia at Augusta University; Charlie Norwood Veterans Affairs Medical Center, Augusta GA 30912

**Keywords:** Drp1, mitochondrial fission, hindlimb ischemia, peripheral arterial disease, macrophage, inflammation, metabolic reprogramming

## Abstract

In the preclinical model of peripheral arterial disease (PAD), M2-like anti-inflammatory macrophage polarization and angiogenesis are required for revascularization. The regulation of cell metabolism and inflammation in macrophages is tightly linked to mitochondrial dynamics. Drp1, a mitochondrial fission protein, has shown context-dependent macrophage phenotypes with both pro- and anti-inflammatory characteristics. However, the role of macrophage Drp1 in reparative neovascularization remains unexplored. Here we show that Drp1 expression was significantly increased in F4/80^+^ macrophages within ischemic muscle at day 3 after hindlimb ischemia (HLI), an animal model of PAD. Myeloid-specific Drp1^-/-^ mice exhibited reduced limb perfusion recovery, angiogenesis and muscle regeneration after HLI. These effects were associated with an increase in pro-inflammatory M1-like macrophages, p-NFkB and TNFα, and reduced anti-inflammatory M2-like macrophages and p-AMPK in ischemic muscle of myeloid Drp1^-/-^ mice. *In vitro*, Drp1^-/-^ macrophages under hypoxia serum starvation (HSS), an in vitro PAD model, demonstrated enhanced glycolysis via reducing p-AMPK as well as mitochondrial dysfunction and excessive mitochondrial ROS, resulting in increased M1-gene and reduced M2-gene expression. Conditioned media from HSS-treated Drp1^-/-^ macrophages exhibited increased secretion of pro-inflammatory cytokines and suppressed angiogenic responses in cultured endothelial cells. Thus, Drp1 deficiency in macrophages under ischemia drives inflammatory metabolic reprogramming and macrophage polarization, thereby limiting revascularization in experimental PAD.

## INTRODUCTION

Peripheral artery disease (PAD) poses a substantial burden of morbidity and mortality due to tissue damage resulting from acute and chronic occlusive ischemia (1, 2). Angiogenesis is the formation of new blood vessels from pre-existing ones, which requires endothelial cells (ECs) to proliferate, migrate, and differentiate to form new vascular structures and is indispensable for restoring limb perfusion of PAD (3). Therapeutic angiogenesis has faced multiple disappointments with clinical trials failing to demonstrate a significant difference in primary endpoint effects (4). Macrophage plays an important role in restoring perfusion recovery and revascularization following ischemia via anti-inflammatory polarization and producing angiogenic factors, which is required for treatment of PAD (5–9). In the early phases of the response to ischemic injury, a hypoxic and pro-inflammatory environment attracts pro-inflammatory M1-type macrophages characterized by enhanced aerobic glycolytic metabolism. Subsequently, a metabolic transition occurs, shifting macrophage phenotypes towards anti-inflammatory M2-type macrophages with increased oxidative phosphorylation in mitochondria, thus promoting reparative angiogenesis and neovascularization (10–13). Macrophage phenotype transition is a complex process influenced by tissue-specific and stimulus-specific differentiation, activation, and maturation programs (14). Defects in normal macrophage metabolism and polarization can lead to unresolved tissue injury, and exacerbation of tissue damage and impaired healing process (10, 11). The critical process of macrophage metabolic reprogramming for tissue repair is in part dependent on mitochondrial metabolism and dynamics (15); however, the underlying mechanisms in the context of ischemia-induced neovascularization are poorly understood.

Mitochondria are highly dynamic organelles that orchestrate metabolic and immune adaptations in response to ischemia. Prolonged ischemia can compromise cellular bioenergetics due to mitochondrial dysfunction (16). Mitochondrial dynamics, continuously adapting to extracellular stimuli, are tightly regulated by the mitochondrial fission proteins including dynamin-related GTPase Drp1, Mitochondrial Fission Factor1 (Mff1) and mitochondrial fusion proteins including Mitofusins (Mfn1and Mfn2) and ortic atrophy1 (OPA1). Recent reports suggest the role of mitochondrial dynamics in regulating mitochondrial reactive oxygen species (mitoROS) levels, calcium homeostasis, and oxidative phosphorylation (17). Altered mitochondrial dynamics in macrophages are a universal response to a variety of inflammatory stimuli. In a context-dependent fashion, Drp1 plays a role in regulating pro- or anti-inflammatory macrophage phenotypes (18–22). It is shown that the inhibition of Drp1 using Mdivi-1 or Drp1 siRNA transfection leads to a reduction in glycolysis, ROS production, and the NFkB-dependent pro-inflammatory M1-macropahge polarization when stimulated with lipopolysaccharide (LPS) (20). By contrast, Drp1 deficiency exacerbates LPS-induced systemic inflammation and liver injury in *in vivo* murine model (19). However, the role of macrophage Drp1 in reparative neovascularization during ischemia in an experimental PAD has never been reported.

In the present study using myeloid-specific Drp1 knockout (*MɸDrp1^KO^*) mice with hindlimb ischemia (HLI) model, an animal model of PAD, we provide the compelling evidence that *MϕDrp1^KO^* mice exhibit impaired blood flow recovery, neovascularization, and muscle regeneration in response to HLI. This was associated with increased pro-inflammatory M1-like macrophage polarization, p-NFkB and TNFα and decreased anti-inflammatory M2-like macrophage polarization and phosphorylation of AMP-activated protein kinase (p-AMPK) in ischemic muscle. *In vitro* studies using hypoxia serum starvation (HSS)-exposed bone marrow-derived macrophage (BMDM) from wild type (WT) and *MɸDrp1^KO^* mice revealed that Drp1^-/-^ BMDM under HSS exhibited enhanced glycolysis via reducing p-AMPK as well as decreased mitochondrial oxygen consumption rate (OCR) and excess mitoROS production, which in turn increased M1-genes and decreased M2-genes, and suppressed angiogenic responses in cultured ECs. Our study will uncover macrophage Drp1 as a positive regulator and therapeutic target for driving reparative neovascularization via promoting anti-inflammatory macrophage polarization and metabolic reprogramming under ischemia.

## Results

### Myeloid Drp1 deficiency impairs reparative neovascularization via inhibiting angiogenesis and arteriogenesis in response to HLI

To assess the role of Drp1 in reparative neovascularization, we used the mouse HLI model which induces ischemia by femoral artery ligation and excision (23). Immunofluorescence analysis revealed that Drp1 was highly expressed in F4/80^+^ macrophages in ischemic muscles vs. non-ischemic muscles at day 3 after HLI (Figure 1A). To determine the role of macrophage Drp1 in post-ischemic neovascularization, we generated mice with myeloid-specific deletion of Drp1 (*MɸDrp1^KO^*) by crossing homozygous *Drp1^fl/fl^* mice with mice expressing Cre recombinase under control of the lysozyme M promoter (LysMCre) (Figure 1B). Selective deletion of Drp1 in macrophage was confirmed by protein analysis in BMDM and peritoneal macrophage, but not in lungs and heart isolated from *MɸDrp1^KO^* and *Drp1^fl/fl^*(control, WT) mice (Figure 1C). Using laser speckle contrast perfusion imaging, we observed that limb perfusion recovery by day 21 post-HLI was significantly reduced in *MɸDrp1^KO^* mice vs. WT mice (Figure 1D). Immunohistochemical analysis showed that the numbers of CD31-positive capillary-like ECs (Figure 1E) as well as α-smooth muscle actin (αSMA)-positive arterioles (Figure S1) in ischemic muscles after HLI were significantly reduced in *MɸDrp1^KO^* mice. H&E staining showed that increase in collateral lumen and wall area in semi-membranous muscle induced by HLI were significantly reduced in *MɸDrp1^KO^* mice (Figure 1F). This was associated with increased necrotic myofibers and delayed muscle regeneration in ischemic muscles of *MɸDrp1^KO^* mice vs. WT mice (Figure 1G). Thus, macrophage Drp1 is required for restoring limb perfusion recovery and neovascularization by promoting angiogenesis and arteriogenesis as well as tissue repair and protection from necrosis in ischemic muscles after HLI.

**Figure 1:**
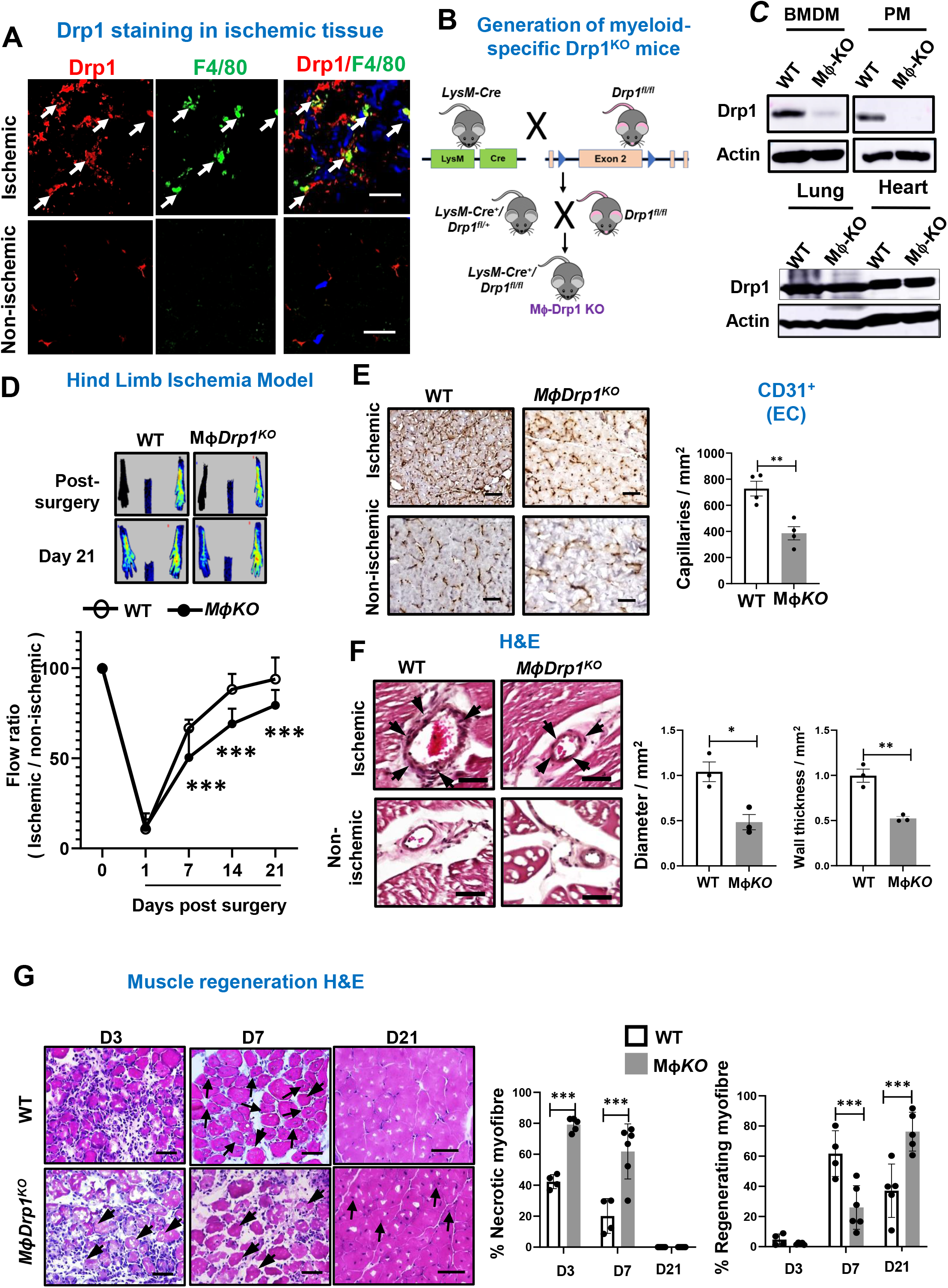
Myeloid Drp1^-/-^ mice exhibited impaired reparative neovascularization via reducing angiogenesis and arteriogenesis in response to ischemia. **A.** Immunofluorescence analysis for Drp1 (red) or F4/80^+^ macrophage (green) expression and their co-localization in non-ischemic and ischemic gastrocnemius (GC) muscles at day 3 post-HLI. Scale bar=10 µm. **B.** Schematic representation of breeding strategy for generating myeloid-specific *Drp1^KO^*(*MɸDrp1^KO^*) mice by crossing LysM-Cre mice with *Drp1^fl/fl^* mice. **C.** Myeloid-specific deletion of Drp1 were shown by immunoblotting (IB) for Drp1 protein expression in bone marrow-derived macrophage (BMDM), peritoneal macrophages (PM), lungs and heart isolated from *Drp1^fl/fl^* (WT) and *MɸDrp1^KO^* mice. **D.** Upper panels show representative laser Doppler images of legs at day 0 and day 21. Lower panels show the blood flow recovery after HLI as determined by the ratio of foot perfusion between ischemic (left) and non-ischemic (right) legs in WT and *MɸDrp1^KO^*mice. **E.** Immunohistochemical analysis for CD31^+^ staining (capillary density) in ischemic and non-ischemic GC muscles in WT and *MɸDrp1^KO^* mice at day21 post-HLI. Scale bar=20µm. The right panel shows quantification. **F**. H&E staining of ischemic and non-ischemic adductor (AD) muscles in WT and *MɸDrp1^KO^*mice at day 7 post-HLI. Arrow heads show collateral arteries. Scale bar=50µm. The right panels show quantification of diameter and wall thickness of collateral arteries. **G**. H&E staining of ischemic and non-ischemic GC muscles in WT and *MɸDrp1^KO^* mice at indicated times after HLI. The right panels show quantification of % of necrotic and regenerating myofiber in these muscles. Scale bar=20µm. Data are mean ± SEM n=4-6. *p<0.05, **\*\***p<0.01, **\*\*\***p<0.001.

### Myeloid Drp1 deficiency increases pro-inflammatory M1-like macrophages and decreases anti-inflammatory M2-like macrophages without affecting the recruitment of Ly6C^hi^ monocytes in ischemic muscles

Macrophages are required for revascularization during HLI (5–9). To examine whether myeloid Drp1 regulates macrophage accumulation and polarization in ischemic hindlimbs, we performed immunofluorescence analysis and found that the numbers of F4/80^+^ macrophages were increased in ischemic muscles of WT mice at days 3 and 7 after HLI, which were significantly reduced in *MɸDrp1^KO^* mice (Figure 2A). This was further confirmed by flow cytometry analysis of isolated F4/80^+^ Mϕ from ischemic muscles of WT and *MɸDrp1^KO^* mice at days 3 and 7 after HLI (Figure 2B). We next examined the polarization of F4/80^+^ macrophages into CD80^+^ pro-inflammatory M1-like macrophages and CD206^+^ anti-inflammatory M2-like macrophages in ischemic muscles. Immunofluorescence analysis showed that the numbers of CD206^+^ M2-like macrophages and CD80^+^ M1-like macrophages in ischemic muscles were increased at days 3 and 7 after HLI in WT mice. But the numbers of CD206^+^ M2-like macrophages was significantly reduced at days 3 and 7 (Figure 2C) while those of CD80^+^ M1-like macrophages were enhanced at day 7 (Figure 2D) in ischemic muscle of *MɸDrp1^KO^* mice vs. WT mice. This is further confirmed by flow cytometry analysis of isolated F4/80^+^ CD206^+^ M2-like macrophages and F4/80^+^CD80^+^ M1-like macrophages in ischemic muscles at days 3 and 7 after HLI (Figure 2E and 2F).

**Figure 2:**
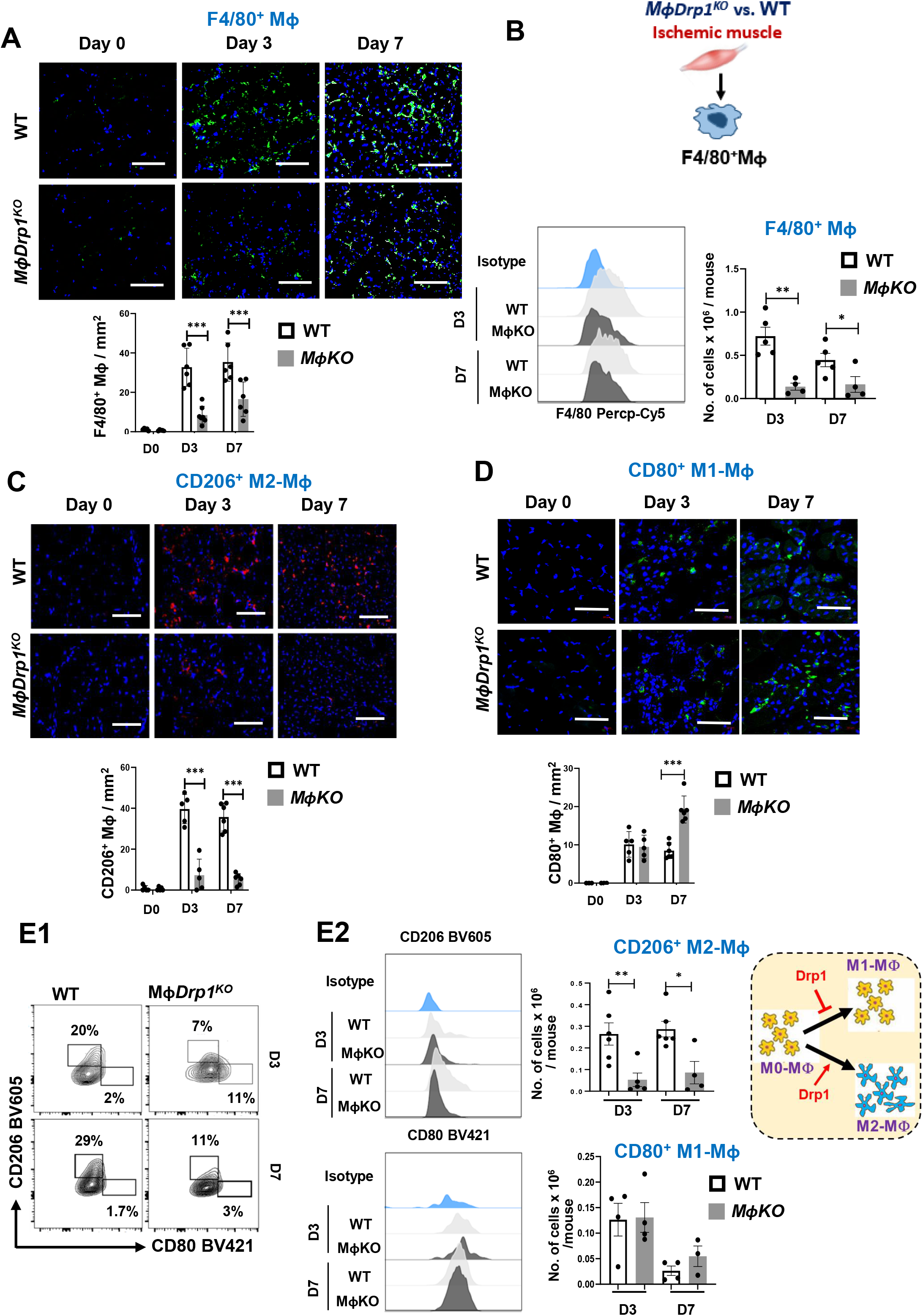
Myeloid Drp1^-/-^ mice exhibited increase in pro-inflammatory M1-like macrophages and decrease in anti-inflammatory M2-like macrophages in ischemic muscles. **A.** Immunofluorescence analysis of F4/80^+^ macrophages (green) with DAPI (blue) in non-ischemic and ischemic GC muscles in WT and *MɸDrp1^KO^*mice at indicated times after HLI. Scale bar=20µm. **B.** Flow cytometry histograms and quantification of numbers of F4/80^+^ macrophages isolated from ischemic muscles in WT and *MɸDrp1^KO^* mice at indicated times after HLI. **C and D**. Immunofluorescence analysis of CD206^+^ M2-like macrophages (red) (**C**) or CD80^+^ macrophages (green) (**D**) with DAPI (blue) in ischemic GC muscles in WT and *MɸDrp1^KO^* mice at indicated times post-HLI. Bottom panels show quantification. **E**. Representative flow cytometry contour plots (E1), and histograms and quantification (E2) of CD45^+^ CD11b^+^ F4/80^+^ CD206^+^ M2-like macrophage and CD45^+^ CD11b^+^ F4/80^+^ CD80^+^ M1-like macrophage numbers in ischemic GC muscles in WT and *MɸDrp1^KO^* mice at indicated time post-HLI. Data are mean ± SEM (n=4-6). *p<0.05, **\*\***p<0.01, **\*\*\***p<0.001.

We next examined whether decrease in F4/80^+^ macrophages in ischemic muscles by myeloid Drp1 deficiency was due to decrease in immune cells mobilization from bone marrow or their recruitment/infiltration or macrophages differentiation. To characterize the immune cells, we performed flow cytometry analysis in bone marrow, peripheral blood and ischemic muscle after HLI (gating strategy and antibody panel are shown in Figure S2, Table 1 & 2). We found that the numbers of Ly6G^+^ neutrophils and Ly6C^hi^ monocytes in peripheral blood (Figure S3A) and bone marrow (Figure S3B) were similar between WT and *MɸDrp1^KO^* mice at days 3 and 7 after HLI. Figure S3C showed that, in ischemic muscles, the numbers of Ly6G^+^ neutrophils were significantly increased in *MɸDrp1^KO^* mice vs. WT mice while those of Ly6C^hi^ monocytes were similar between *MɸDrp1^KO^* mice and WT mice at day 3 after HLI. Then, both Ly6G^+^ neutrophils and Ly6C^hi^ monocytes were reduced by day 7. These results suggest that loss of myeloid Drp1 limits anti-inflammatory macrophage polarization and differentiation of Ly6C^hi^ monocytes to mature macrophages without affecting the releasing monocytes from bone marrow and their recruitment into ischemic muscle, which impairs ischemia-induced revascularization during HLI.

**Table 1:** Enlisted sequence of primers used in q-RT-PCR analysis.

**Table 2:** Enlisted catalog of antibodies used for flow cytometric and immunofluorescence analysis.

### Myeloid Drp1 deficiency increased pro-inflammatory genes and signaling and decreased anti-inflammatory genes and signaling in ischemic muscle

To confirm further the role of myeloid Drp1 in regulating macrophage polarization during HLI, we isolated F4/80^+^ macrophages from ischemic muscle at day 3 after HLI and found that pro-inflammatory M1-marker genes including *Nos2*, *Ptgs2* and *Il-1β* were up-regulated while anti-inflammatory M2-marker genes including *Arg1* and *Retnla* were downregulated in *MɸDrp1^KO^* mice vs. WT mice (Figures 3A). This was associated with significant decrease in pro-angiogenic growth factor genes such as *Vegf* and *Tgfβ* in macrophages isolated from ischemic muscles of *MɸDrp1^KO^* mice vs. WT mice (Figure 3A).

**Figure 3:**
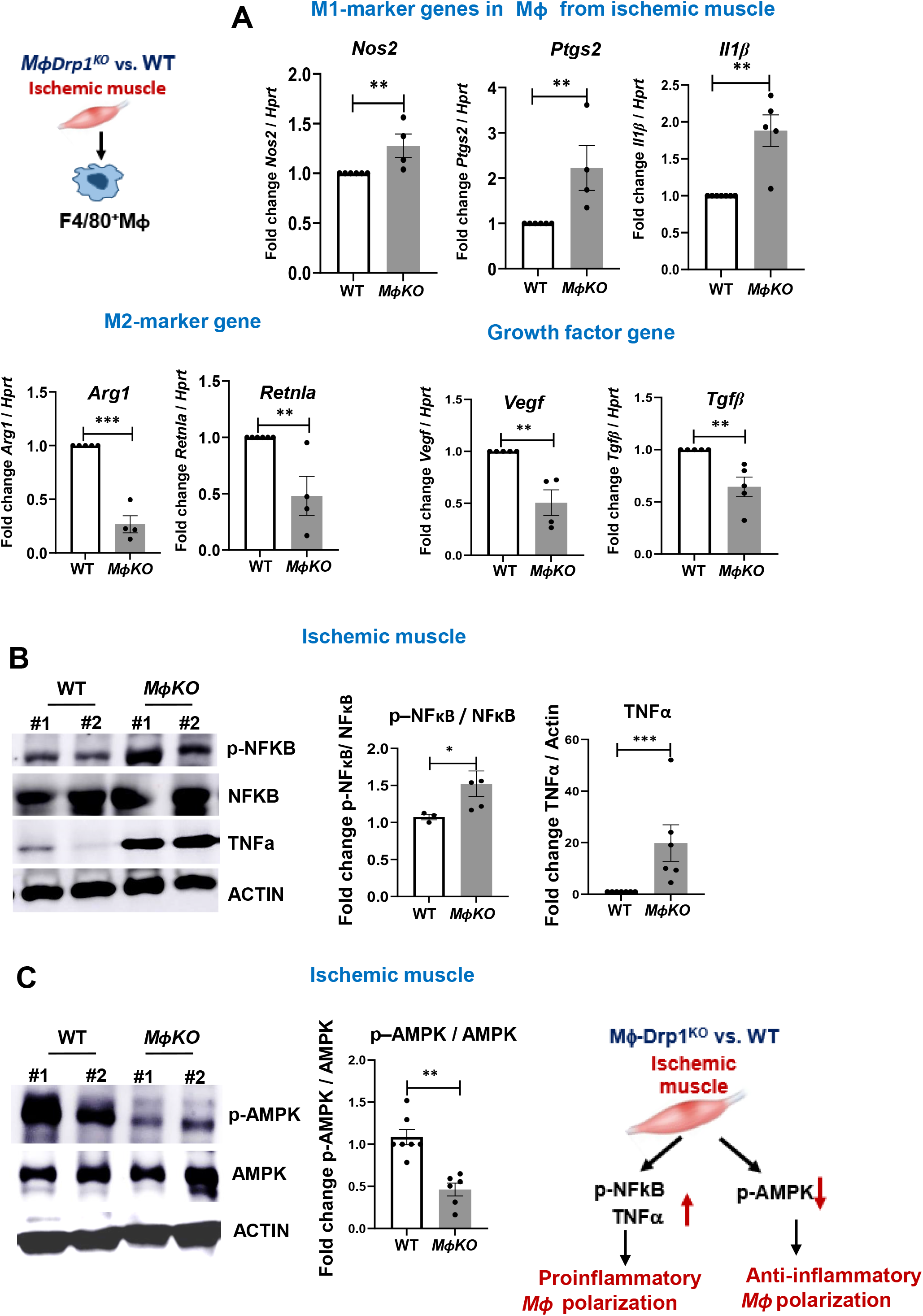
Myeloid Drp1^-/-^ mice exhibited increase in pro-inflammatory genes and signaling and decrease in anti-inflammatory genes and signaling in ischemic muscle. **A.** Left, schematic representation of magnetic bead based F4/80^+^ macrophage purification from ischemic GC muscle at day 3 post-HLI. These macrophages were used to analyze the pro-inflammatory genes (*Nos2*, *Ptgs2* and *Il1β*), anti-inflammatory genes (*Arg1* and *Retnla*) and pro-angiogenic growth factor genes (*Vegf* and *Tgfβ*) mRNAs using quantitative (q)RT-PCR (fold increase relative to *Hprt*) in WT and *MɸDrp1^KO^* mice at day 3 post-HLI. **B and C.** Western blot analysis for the pro-inflammatory p-NFκB, NFκB and TNFα protein (B) as well as anti-inflammatory p-AMPK and AMPK protein (C) in ischemic tibialis anterior (TA) muscles in WT and *MɸDrp1^KO^* mice at day 3 post-HLI. The right panels show the quantification. Data are mean ± SEM. *p<0.05, **p<0.01, ***p<0.001.

Previous studies showed that pro-inflammatory NFκB signaling and anti-inflammatory AMPK signaling contribute to macrophage polarization toward M1- and M2-phenotype, respectively (24–27). We found that expression of p-NFκB, but not total NFkB protein, was significantly increased in ischemic muscles on day 3 post-HLI of *MɸDrp1^KO^* mice vs. WT mice, which was associated with increased expression of pro-inflammatory TNFα (Figure 3B). In contrast, expression of p-AMPK, but not total AMPK protein, was significantly reduced in ischemic tissues on day 3 post-HLI of *MɸDrp1^KO^* mice vs. WT mice (Figure 3C). Taken together, these results suggest that myeloid Drp1 deficiency increased pro-inflammatory genes and signaling (p-NFkB)-M1-like macrophage axis, while decreased anti-inflammatory genes and signaling (p-AMPK)-M2-like polarization of macrophage axis in ischemic muscles, which resulted in impaired reparative neovascularization after HLI.

### Drp1^-/-^ BMDM under HSS induced mitochondrial hyperfusion, which was associated with increased M1-macrophage and decreased M2-macrophage polarization via reducing p-AMPK

To address the underlying mechanism by which Drp1 deficiency regulates macrophage polarization under ischemia, we performed in vitro studies using BMDM isolated and cultured from WT and *MɸDrp1^KO^* mice under hypoxia serum starvation (HSS), an *in vitro* PAD model (Figure 4A), as previously described (28, 29). Mito-tracker staining showed that WT-BMDM under HSS induced mitochondrial fission, which peaked at 1h and 2h and returned to basal level within 8h (Figure S4A). However, HSS did not significantly alter pSer616-Drp1 or pSer637-Drp1 or expression of Drp1 protein or mitochondrial fission protein Mff1 or mitochondrial fusion proteins Mfn1, Mfn2 and OPA1 in BMDM (Figure S4B). In contrast, Drp1^-/-^ BMDM showed elongated mitochondrial fusion at 2h and 8h (not shown) after HSS stimulation (Figure 4B). We next examined the role of Drp1 in macrophage polarization using BMDM under HSS *in vitro*. Consistent with *in vivo* HLI model, HSS exposure to BMDM increased M1 marker gene *Nos2* and decreased M2 marker gene *Retnla* in a time-dependent manner (Figure S5A). Drp1^-/-^ BMDM with HSS exhibited enhanced M1-marker genes (*Nos2* and *Ptgs2*) and further decreased M2-maker genes (*Arg1* and *Retnla*) vs. WT BMDM with HSS (Figure S5B). These findings suggest that loss of Drp1 in macrophage under HSS promotes pro-inflammatory macrophage polarization and reduces anti-inflammatory macrophage polarization *in vitro*.

**Figure 4:**
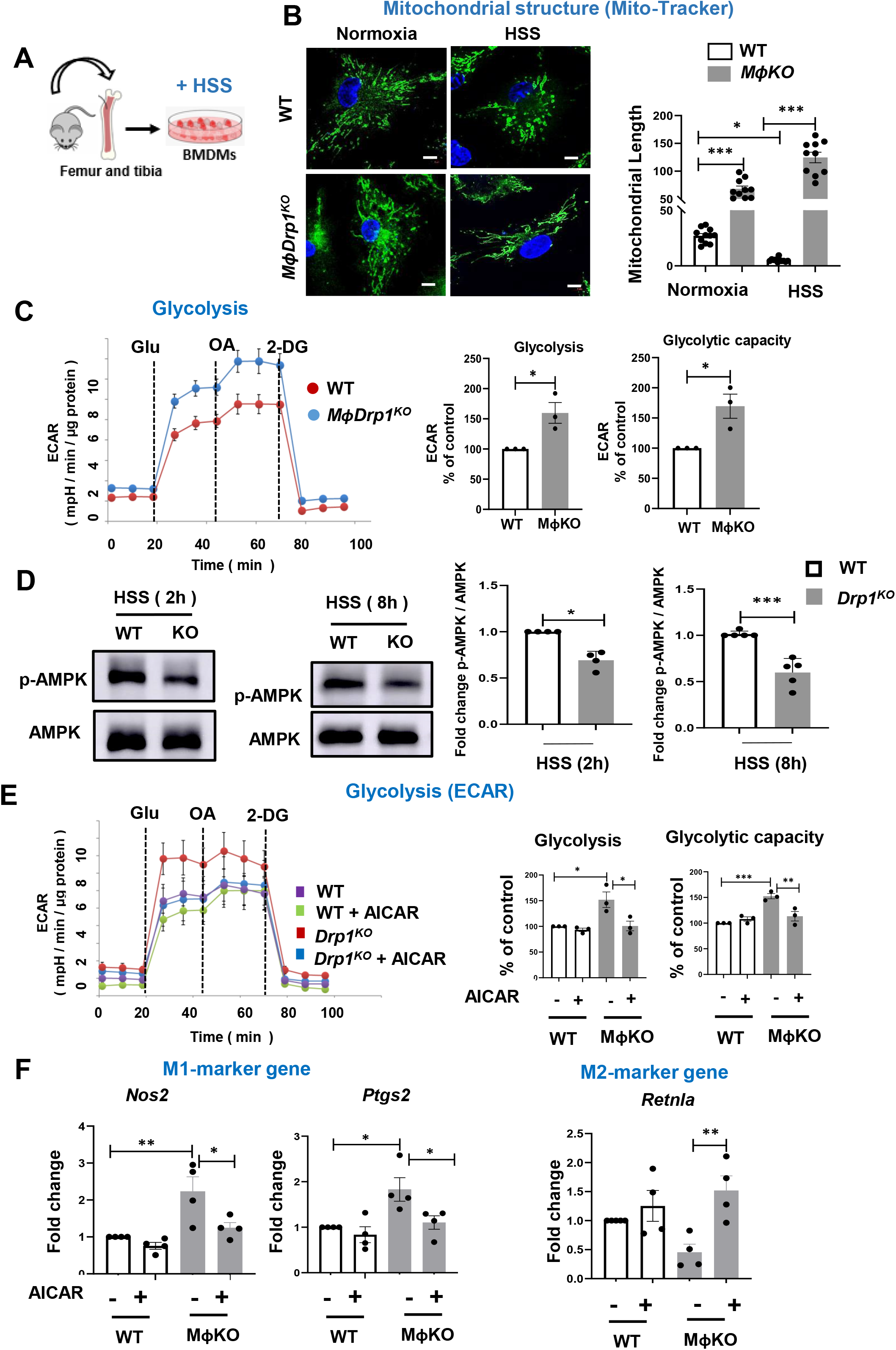
Drp1^-/-^ BMDM under HSS exhibited mitochondrial hyperfusion and hyperglycolysis, which was associated with decreased M2-like macrophages via reducing p-AMPK. **A.** Schematic representation showing cultured mouse BMDM exposed to hypoxia serum starvation (HSS), an *in vitro* PAD model. BMDM from WT and *MɸDrp1^KO^*mice were cultured for 7 days and then exposed to HSS. **B.** Analysis of mitochondrial structure by Mito-tracker green staining in WT and Drp1^-/-^BMDM with or without HSS stimulation for 2 h. Right panel shows quantification of mitochondrial length by Image J. Scale bar=5μm. **C.** Analysis of glycolysis measured by extracellular acidification rate (ECAR) using Seahorse XF analyzer in WT and Drp1^-/-^BMDM under HSS for 2 h. Right panels show quantification. n=3. **D.** Western blot analysis for p-AMPK and AMPK expression in WT and Drp1^-/-^BMDM with HSS stimulation for 2 h and 8 h. Right panels show quantification. n=3-5. **E and F**. Effects of AMPK activator AICAR (pre-treatment for 2 h at 100 μM) on glycolysis (ECAR) measured by Seahorse XF analyzer (E) as well as mRNA expression for M1-markers (*Nos2* and *Ptgs2*) and M2-marker *Retnla* measured by qRT-PCR (F) in WT and Drp1^-/-^BMDM under HSS for 2 h (E) or 8 h (F). Data are mean ± SEM. n=3-4. *p<0.05, **p<0.01, ***p<0.001.

Since pro-inflammatory macrophage polarization is shown to undergo metabolic reprograming from oxidative phosphorylation to aerobic glycolysis (12, 13), we measured the extracellular acidification rate (ECAR) using Seahorse assay. Figure 4C showed that Drp1^-/-^ vs. WT BMDM under HSS exhibited increased glycolysis and glycolytic capacity. To address the mechanism by which Drp1 deficiency in macrophage under HSS enhances glycolysis, we measured p-AMPK which has been shown to promote anti-inflammatory and pro-angiogenic M2-Mɸ polarization via suppression of glycolysis (26, 27). Consistent with ischemic tissue in vivo, we found that p-AMPK was significantly decreased in Drp1^-/-^ BMDM vs. WT BMDM under HSS (Figure 4D). To determine whether decreased p-AMPK contributes to enhanced glycolysis and pro-inflammatory macrophage polarization in Drp1^-/-^ BMDM under HSS, we performed rescue experiments using AMPK activator 5-aminoimidazole-4-carboxiamide ribonucleotide (AICAR) (27, 30). We found that the activation of AMPK by AICAR, at a concentration that did not alter glycolysis in WT BMDM subjected to HSS, effectively restored augmented glycolytic activity in Drp1^-/-^ BMDM under HSS (Figure 4E). Furthermore, this AMPK activation restored the elevated expression of pro-inflammatory M1-genes *Nos2* and *Ptgs2* in Drp1^-/-^ BMDM under HSS as comparable to those observed in HSS-exposed WT BMDM (Figure 4F). Notably, AICAR also rescued the reduced expression of genes associated with the anti-inflammatory M2 phenotype in Drp1^-/-^ BMDM under HSS (Figure 4F).

### Drp1^-/-^ BMDM under HSS induced mitochondrial dysfunction and mitoROS production, which contributes to promoting M1-like macrophage polarization and inhibiting M2-like macrophage polarization

We next examined the role of macrophage Drp1 in mitochondrial respiration by measuring OCR using Seahorse assay. Figure 5A showed that Drp1^-/-^ BMDM under HSS exhibited a significant decrease in basal respiration, maximal respiration, and ATP production vs. WT BMDM with HSS. This mitochondrial dysfunction was associated with excess mitoROS production in Drp1^-/-^ BMDM under HSS, as measured by MitoSOX fluorescence (Figure 5B). We then investigated the impact of elevated mitoROS due to a deficiency in Drp1 on *Mɸ* polarization under ischemic conditions. Our results demonstrate that treatment with the mitochondria-specific O_2_^-^ scavenger, Mito-TEMPO, effectively rescued the increase in pro-inflammatory M1-genes *Nos2* and *Ptgs2*, as well as the decrease in the M2-marker gene *Retnla*, in Drp1^-/-^ BMDM exposed to HSS. Notably, this treatment did not affect the response of WT-BMDM under HSS conditions (Figure 5D). These findings strongly imply that myeloid Drp1 deficiency leads to mitochondrial dysfunction and an excessive production of mitoROS, resulting in an augmented M1-like macrophage polarization and compromised M2-like macrophage polarization in an *in vitro* ischemic condition. It is important to note that the application of Mito-TEMPO had no significant impact on the increased glycolysis observed in Drp1^-/-^ BMDM under HSS, as shown in Figure 5C. Additionally, we observed that the activation of AMPK by AICAR did not mitigate the excess mitoROS production in Drp1^-/-^ BMDM under HSS (Figure S6A). Consequently, our results suggest that the concurrent elevation of excess mitoROS and glycolysis coupled with reduced AMPK activation, collectively contribute to the promotion of M1-like macrophage polarization and the inhibition of M2-like macrophage polarization in Drp1^-/-^ BMDM under HSS conditions.

**Figure 5:**
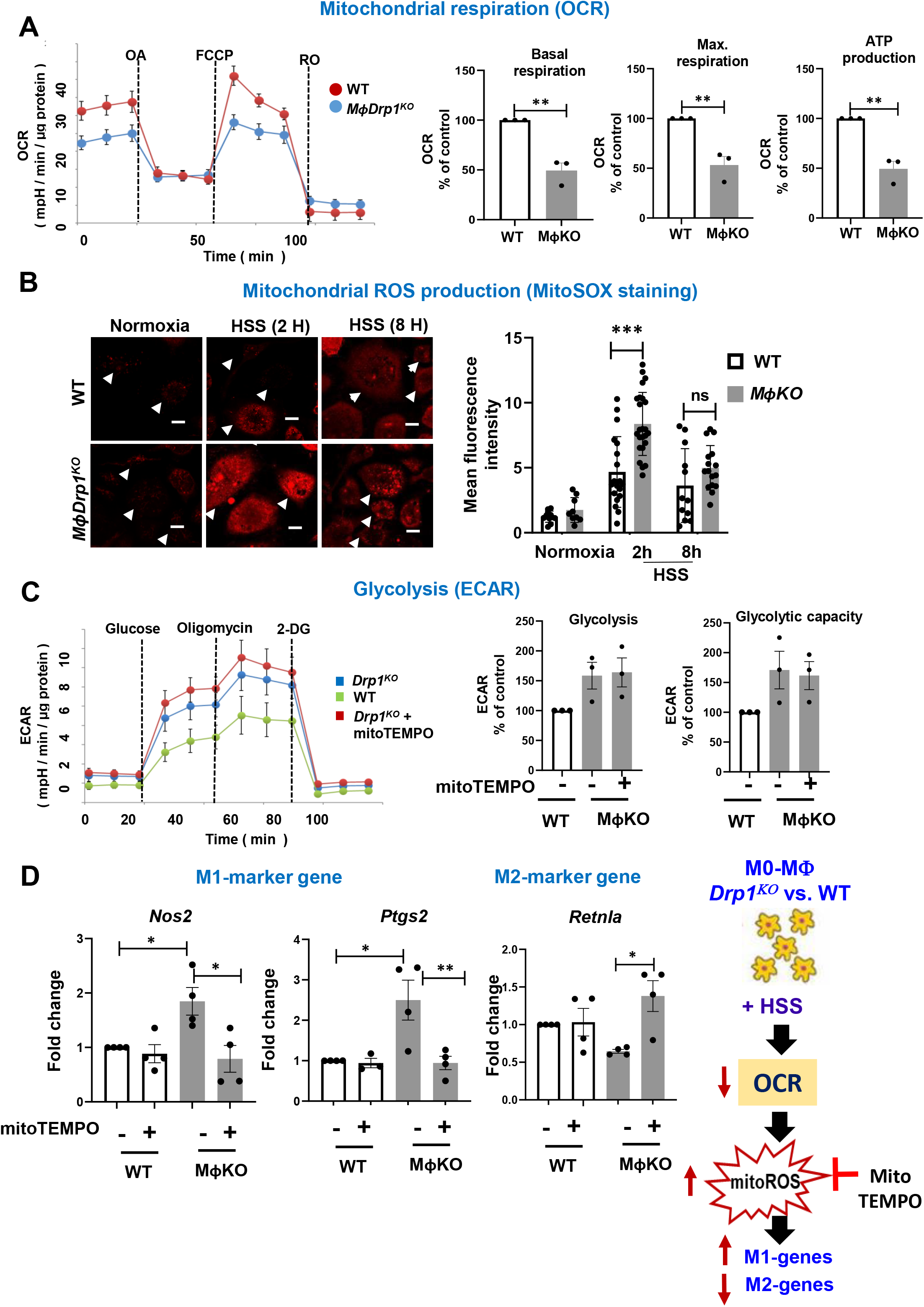
Drp1^-/-^ BMDM under HSS induced mitochondrial dysfunction and mitochondrial ROS (mitoROS) production, which in turn promoted M1-like macrophage polarization and reduced M2-like macrophage polarization. **A.** Analysis of mitochondrial respiration measured by OCR using Seahorse XF analyzer in WT and Drp1^-/-^BMDM under HSS for 2 h. Right panels show quantification. **B.** Mitochondrial ROS production measured by MitoSOX under normoxia or HSS for 2 h and 8 h in WT and Drp1^-/-^ BMDM. Right panels show the averaged fluorescence intensity quantified by Image J. Scale bars=5 μm. **C and D**. Effects of mito-TEMPO (pre-treatment of 16 h at 20 μM) on glycolysis (ECAR) using Seahorse assay (C) as well as mRNA expression for M1-markers (*Nos2* and *Ptgs2*) and M2-marker *Retnla* measured by qRT-PCR (D) in WT and Drp1^-/-^BMDM under HSS for 2 h (C) or 8 h (D). Data are mean ± SEM. n=4. *p<0.05, **p<0.01, ***p<0.001.

### Conditioned media from WT-BMDM under HSS promotes angiogenesis in ECs, which is inhibited by those from Drp1^-/-^ BMDM under HSS

To further elucidate the influence of myeloid Drp1 on the regulation of angiogenesis in ECs, we first collected and analyzed the effect of paracrine factors present in the conditioned media released by WT and Drp1^-/-^ BMDM exposed to HSS (Figure 6A). The conditioned media obtained from HSS-exposed WT-BMDM markedly promoted the migration of EC in response to a wound scratch assay on confluent EC cultures (Figure 6B). These pro-angiogenic effects were absent when conditioned media from HSS-treated *Drp1^KO^*BMDM were employed. This was associated with the observation that the conditioned media from Drp1^-/-^ BMDM lead to an increased secretion of pro-inflammatory cytokines, including TNFα and IL-6 (Figure 6C). These results are in accordance with our *in vivo* observations and underscore the important role of macrophage Drp1 in mediating the crosstalk between macrophage and ECs to facilitate ischemia-induced angiogenesis.

**Figure 6:**
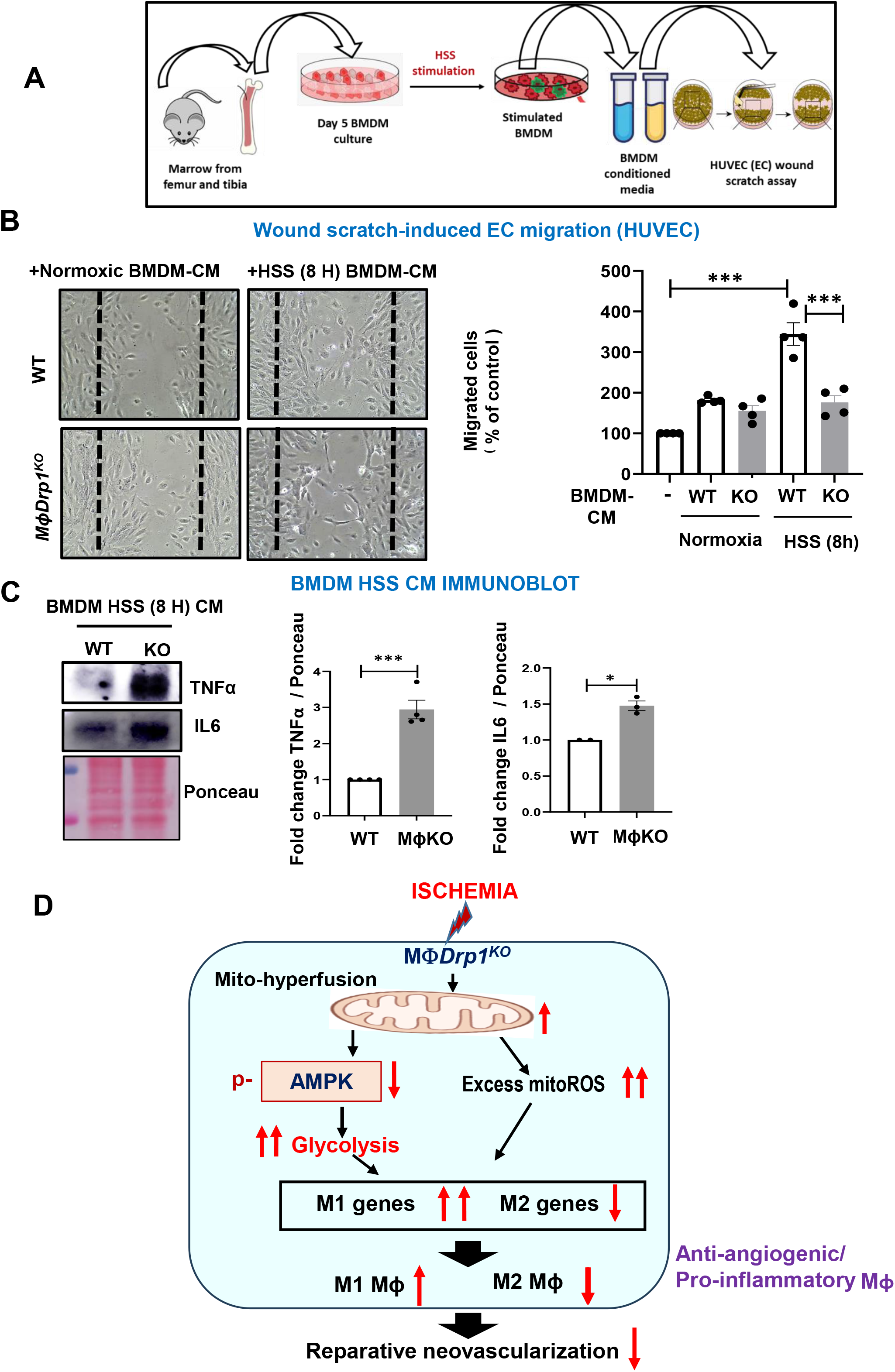
Conditioned media from WT-BMDM under HSS promoted angiogenesis in ECs, which was inhibited by those from Drp1^-/-^ BMDM under HSS. **A.** Schematic representation of HSS-stimulated mouse BMDM conditioned media (CM) collection and their treatment on confluent HUVEC upon wound scratch. **B.** Analysis of EC migration using wound-scratch assay in confluent monolayers of HUVEC followed by treatment with normoxic control or HSS-BMDM CM (20 ng/mL). Representative bright field images (left) and quantification of number of migrated cells per field (right) in HUVECs at 0h and 16h after wound scratch. Scale bars=20 μm. **C**. Protein expression of pro-inflammatory TNFα and IL-6 in WT and Drp1^-/-^BMDM CM under HSS for 8 h. Right panels show quantification. n=3. Data are mean ± SEM. *p<0.05, **p<0.01, ***p<0.001. **D**. Schematic model showing that Drp1 deficiency in macrophages under ischemia induces metabolic reprograming (hyperglycolysis) via reducing AMPK activation as well as mitochondrial dysfunction-excess mitoROS axis, thereby increasing pro-inflammatory M1-like macrophage polarization and reducing anti-inflammatory M2-like macrophage polarization, which, in turn, limits reparative neovascularization in response to ischemic injury.

## DISCUSSION

Macrophages play a pivotal role in the process of restoring perfusion recovery and revascularization following ischemia primarily through their capacity for anti-inflammatory M2-polarization and the production of angiogenic factors. Their functions are indispensable for the effective treatment of PAD (5–9). In the present study, we demonstrate previously unidentified roles of myeloid Drp1 as a key regulator of reparative macrophage responses during ischemia, with implications for therapeutic neovascularization following ischemic injury. This regulation is mediated through promoting anti-inflammatory macrophage polarization and metabolic reprogramming. Here we show that: 1) Myeloid-specific Drp1^-/-^ mice showed impaired limb perfusion, angiogenesis, arteriogenesis, and muscle regeneration in ischemic muscle after HLI. These effects are concomitant with promoting pro-inflammatory M1-like macrophage polarization, p-NFkB, elevated TNFα levels and a decrease in anti-inflammatory M2-like macrophage polarization and p-AMPK in the ischemic muscle microenvironment; 2) *In vitro*, Drp1^-/-^ BMDM exposed to HSS, an in vitro model of PAD, exhibit enhanced glycolytic activity via a reduction in p-AMPK as well as mitochondrial dysfunction and excess mitoROS production, which leads to an increase in M1-gene expression and a decrease in M2-gene expression; 3) Conditioned media derived from HSS-treated Drp1^-/-^ BMDM reveal an augmented secretion of pro-inflammatory cytokines, and concurrent suppression of angiogenic responses in cultured ECs.

Accumulating evidence suggest the critical role of mitochondrial dynamics as a central intracellular signaling platform for the regulation of innate immune responses (17, 31, 32). However, the specific function of macrophage Drp1 in the context of the inflammatory response remains a subject of conflicting outcomes. For instance, emerging evidence points to reduced inflammatory responses in macrophages and microglia in the absence of Drp1 (18, 21). Furthermore, *in vivo* pharmacological inhibition of Drp1 using the small molecule inhibitor Mdivi-1 has been shown to suppress inflammatory cytokine production and impede inflammation in the context of infections and inflammatory diseases (20, 33, 34). Notably, the impact of myeloid Drp1 varies in different pathological scenarios. For example, macrophage Drp1 promotes vascular injury-induced intimal thickening by enhancing pro-inflammatory macrophage responses (18). Conversely, a deficiency of liver-specific Drp1 exacerbates *in vivo* LPS-induced acute liver injury through an augmented inflammatory cytokine response (19). *In vitro* studies using Drp1 knockdown macrophages have demonstrated enhanced mitochondrial fusion and increased IL-1β production due to activation of the NLRP3 inflammasome (22). Hence, the role of the mitochondrial fission protein, Drp1 GTPase, in regulating pro- or anti-inflammatory phenotypes appears to be contingent on the specific context and cell type. However, the role of macrophage Drp1 in the context of ischemia-induced neovascularization with an experimental model of PAD has remained unknown.

In our present study, we observed that mice with myeloid Drp1 deficiency exhibited impaired blood flow recovery and angiogenesis/arteriogenesis in ischemic limbs, which led to a reduced capacity for muscle regeneration following HLI. These findings suggest an endogenous protective role for myeloid Drp1 in the reparative neovascularization in response to tissue ischemia. Flow cytometry and immunofluorescence analysis revealed that *MɸDrp1^KO^* mice displayed a notable reduction in naïve M0-like F4/80^+^ macrophages and CD206^+^ M2-like macrophages on days 3 and 7 post-HLI, without a significant impact on the recruitment of Ly6C^hi^ monocytes into the ischemic muscles. Of note, *MɸDrp1^KO^* mice exhibited higher numbers of neutrophils in the ischemic muscle on day 3 post HLI, followed by an increase in CD80^+^ M1-like macrophages and reduction of M2-like macrophages on day 7 after HLI. It is shown that Drp1 is involved in the efferocytosis of apoptotic neutrophils, an action that contributes to M2-like anti-inflammatory macrophage polarization (35). Thus, these findings suggest that, in an ischemic context, macrophage Drp1 plays a crucial role in facilitating monocyte to macrophage differentiation and the promotion of M2-like anti-inflammatory macrophage polarization with the inhibition of pro-inflammatory macrophage polarization.

Our present study found that macrophage Drp1 deficiency-driven impaired inflammatory M2-like macrophage phenotype in ischemic muscles is likely due to increasing the pro-inflammatory NFkB signaling and a concurrent decrease in the activation of the anti-inflammatory p-AMPK. This finding aligns with previous reports demonstrating the detrimental role of pro-inflammatory NFkB in hypoxia-induced ischemic brain injury and inflammation (24, 25). In contrast, AMPKα1 has been shown to promote HLI-induced arteriogenesis by regulating the production of angiogenic growth factors such as VEGF, TGFβ, and FGF2 by M2-like macrophages (26, 27). It’s important to note that pro-inflammatory macrophages shift their metabolic profile from oxidative phosphorylation to aerobic glycolysis (12, 13). AMPK has been shown to promote anti-inflammatory and pro-angiogenic M2-macrophage polarization by suppressing pro-inflammatory glycolysis (26, 27). In line with our in vivo observations in ischemic tissue, we found a notable reduction in p-AMPK in Drp1^-/-^ BMDM when compared to WT-BMDM in our *in vitro* studies. Under HSS, loss of Drp1 led to the enhanced induction of M1 genes *Nos2* and *Ptgs2* through the reduced activation of AMPK in BMDM. Consequently, conditioned media from myeloid Drp1 KO macrophages exposed to HSS exhibited the secretion of pro-inflammatory cytokines TNFα and IL-6 and had impaired pro-angiogenic effects of WT macrophages on cultured ECs. Mechanistically, the decreased p-AMPK in Drp1^-/-^ BMDM was associated with enhanced glycolysis, as evidenced by the ECAR, an effect that was restored by the treatment with AMPK activator AICAR. AMPK activation also reduced the elevated levels of M1-genes *Nos2* and *Ptgs2* in Drp1^-/-^ BMDM under ischemic conditions, while increasing the lowered levels of M2-gene *Retnla*.

It remains unknown how AMPK activation reduces glycolysis and pro-inflammatory macrophage polarization. It is reported that several downstream targets of AMPK suppress pro-inflammatory signaling in macrophages. These include the activation of PI3 kinase or SIRT1, leading to the deacetylation and inhibition of NFkB activity (26, 36). Additionally, the anti-inflammatory IL-10-mediated rapid activation of AMPK in macrophages is essential for the activation of the PI3K/Akt/mTORC1 and STAT3-mediated anti-inflammatory polarization of macrophages (26, 37). AMPK also is shown to downregulate inflammatory metabolism by inhibiting HIF1α through the inactivation of mTORC1 (38). Further investigations are needed to elucidate the downstream targets of AMPK under ischemic conditions, which drive anti-inflammatory signaling in macrophages.

Mitochondrial functions are crucial for cellular metabolism under ischemic conditions, and prolonged ischemia exacerbates mitochondrial dysfunction (16). While the role of macrophage Drp1 in mediating pro-inflammatory responses in the context of bacterial LPS stimulation is well-documented (20, 34), its involvement in mitochondrial metabolism and inflammatory responses following ischemia remains uncertain. For instance, in response to LPS stimulation, the activation of Drp1 in macrophages leads to the production of excessive mitoROS, which, in turn, further amplifies the secretion of inflammatory cytokines through the activation of NFkB in a Drp1-dependent manner (20). In contrast, we observed that, under ischemic conditions, Drp1^-/-^ BMDM exposed to HSS, an in vitro PAD model, exhibited mitochondrial dysfunction, as evidenced by a decrease in OCR and an excess mitoROS production. Consequently, this led to an upregulation of pro-inflammatory M1-associated genes and a downregulation of an anti-inflammatory M2-associated genes in BMDM. Importantly, these responses in Drp1-deficient macrophages were mitigated by the mitoROS scavenger Mito-TEMPO. Of note, the role of mitoROS in AMPK function has been reported (39, 40). For instance, AMPK-mediated Drp1 regulation prevents EC dysfunction by suppressing mitoROS and ER stress (40). In hypoxic conditions, mitoROS can directly activate AMPK through phosphorylation, independent of the AMP/ATP ratio (39). However, in the present study under ischemic conditions, AICAR treatment did not significantly reduce the enhanced mitoROS levels in Drp1^-/-^ BMDM. Thus, our results suggest that the increased glycolysis, resulting from reduced AMPK activation under HSS in Drp1^-/-^ BMDM, promotes M1-like macrophage polarization independently of an increase in mitoROS.

In our *in vivo* experiments using a preclinical PAD model, we found that myeloid-specific Drp1^-/-^ mice exhibited impaired angiogenesis with enhanced M1- and reduced M2-macrophage polarization in ischemic muscles. Mechanistically, conditioned media obtained from HSS-treated Drp1^-/-^ BMDM revealed loss of pro-angiogenic function of conditioned media from HSS-treated BMDM, which was associated with an augmented secretion of pro-inflammatory cytokines, including TNFα and IL-6. Collectively, our findings suggest that Drp1 deficiency in macrophage under ischemic conditions promotes inflammatory macrophage polarization, partially through the excessive production of mitoROS, which in turn limits revascularization.

In summary, our findings underscore the essential role of macrophage Drp1-mediated mitochondrial dynamics in driving metabolic reprogramming towards anti-inflammatory macrophages, thereby enhancing neovascularization in response to ischemia. Specially, we have identified the activation of AMPK and the concurrent reduction in excessive mitoROS production as crucial downstream targets through which macrophage Drp1 orchestrates the metabolic reprogramming and mitochondrial functions of macrophages, fostering enhanced neovascularization and tissue repair in response to ischemic injury. Furthermore, given the promising results of clinical trials involving intramuscular cell therapy with bone marrow mononuclear cells in patients with critical limb ischemia (4), our findings uncover macrophage Drp1 as a novel and promising therapeutic target for the treatment of PAD.

## Materials and Methods

### Ethics Statement for Animal Study

All animal studies were carried out following protocols approved by the institutional Animal Care Committee and institutional Biosafety Committee at Augusta University. Room temperature and humidity were maintained at 22.5 °C and between 50% and 60%, respectively. All mice were held under the 12:12 (12-h light: 12-h dark) light/dark cycle. Mice were held in individually ventilated caging with a maximum of 5 or a minimum of 2 mice per cage. *Drp1^fl/fl^* (control, wild type (WT)) and myeloid-specific Drp1^-/-^ (*MɸDrp1^KO^*) mice were used at 8-12 weeks (for hindlimb ischemia model). *MɸDrp1^KO^* mice were generated by crossing *Drp1^fl/fl^* mice with *LysM-CRE^Tg/Tg^*mice on a C57BL/6J background. Male mice heterozygous for Drp1 floxed allele and LysM-Cre transgene (Drp1^fl/+^ LysM-Cre^Tg/+^) were crossed-bred with homozygous Drp1 floxed female mice. Both male and female mice with genotype *Drp1^fl/fl^ LysM-Cre^Tg/+^* and the littermates with genotype *Drp1^fl/fl^ LysM-Cre^-/-^* were used at 8-12 weeks.

### Hindlimb ischemia model

Mice were subjected to unilateral hindlimb surgery under anesthesia with intraperitoneal administration of ketamine (87 mg/kg) and xylazine (13 mg/kg). We performed ligation and segmental resection of left femoral artery. Briefly, the left femoral artery was exposed, ligated both proximally and distally using 6-0 silk sutures and the vessels between the ligatures were excised without damaging the femoral nerve. Skin closure was done using 6-0 nylon sutures. We measured ischemic (left)/non-ischemic (right) limb blood flow ratio using a laser Doppler blood flow (LDBF) analyzer (PeriScan PIM 3 System; Perimed) as we described (41, 42). Mice were anesthetized and placed on a heating plate at 37°C for 10 minutes to minimize temperature variation. Before and after surgery, LDBF analysis was performed in the plantar sole. Blood flow was displayed as changes in the laser frequency, represented by different color pixels, and mean LDBF values were expressed as the ratio of ischemic to non-ischemic LDBF.

### Histological analysis

For cryosections, mice were euthanized and perfused through the left ventricle with saline, limbs were fixed in 4% paraformaldehyde (PFA) overnight and incubated with 30% sucrose, and gastrocnemius muscles were embedded in OCT compound (Sakura Finetek). 7 µm cryosections for capillary density were stained with anti-mouse CD31 antibody (MEC 13.3, BD Bioscience). For immunohistochemistry, we used R.T.U. Vectorstain Elite (Vector Laboratories) followed by DAB visualization (Vector Laboratories). Arterioles were stained with Cy3-conjugated anti-αSMA antibody (1A4, Sigma). Macrophages were labeled with anti-F4/80 (BM8, BiolEgend), M1-macrophage were labeled with anti-CD80 (16-10A1, Biolegend) and M2 macrophages were labeled with anti-CD206 (C068C2, Biolegend) antibody. Images were captured by Keyence microscope (bz-x800) or confocal microscopy (Zeiss) and analyzed by Image J or LSM510 software (Zeiss), respectively.

### Isolation and primary culture of bone marrow derived macrophages (BMBMs)

BMDMs were harvested from hind leg tibiae and femur of WT and *MɸDrp1^KO^* mice. Briefly, bone marrow cells were flushed from the bone with a 27G needle in DPBS and filtered using 70 µm filter. Cells were cultured and differentiated in DMEM medium (Gibco) supplemented with antibiotics, 10% FBS and 20% conditioned media from L929 cell line (enriched in CSF-1) for 7 days in polystyrene culture plates and non-adherent cells were washed and removed every alternate day. The resulting BMDM population was determined by staining with anti-CD11b anti-Ly6C and anti-F4/80 (antibody catalog number are mentioned in supplementary Table S2) and assessed by flow cytometry.

### In Vitro Hypoxia Serum Starvation (HSS)

Day 7 BMDMs were washed twice to remove traces of FCS and then incubated in starvation medium from Cell applications Inc. (Cat No: 209-250) and subjected to hypoxia (2% O_2_) for indicated times (28, 29)

### HSS stimulated BMDM conditioned media preparation

BMDM were subjected to HSS for 8 h. Later macrophage conditioned-medium (CM) was collected and passed through 0.2 um filters and stored at -80°C until further use. Just before the specified experiment. BMDM CM were thawed in presence of protease and peptidase inhibitor and concentrated using 10K Amicon ultra centrifugal tubes (UFC801024). Brifely, CM were spun at 4200 RPM for 8 min at 4°C. Total proteins were estimated using Bradford Assay.

### Flow cytometry (FACS) analysis

Mouse whole blood and bone marrow were collected by cardiac puncture and from femur and tibia respectively using 27G needle in EDTA coated 1mL syringe and placed in 1.5 mL EDTA coated Eppendorf tube for 30 min at room temperature. The peripheral blood was spun down at 500 g for 1 min at 4°C. The supernatant was discarded, and the cells were re-suspended in 1 ml of RBC lysis buffer (00-4300-54, Invitrogen) for 10 min at room temperature in dark. Samples were then washed with DPBS with 2% FCS and centrifuged at 500 g for 5 min at 4°C. Cells were re-suspended in cold DPBS with 2% FCS and placed on ice. Excised gastrocnemius muscles were minced and enzymatically digested in DPBS containing collagenase type I 1 mg/mL Type I (C0130, Sigma), 18 mg/mL collagenase Type XI (C7657, Sigma), 1 mg/mL hyaluronidase (H3506, Sigma) and 50 U/mL DNAse (D4263, Sigma) at 37°C for 1 hour. Cells were spun and re-susupended in DPBS with 2% FCS followed by passing through 70 µm and 40 µm filter and counted. The cells suspension was incubated with blocking buffer consisting of anti-mouse CD16/CD32 (14-0161-85, eBioscince) and 2% FCS for 15 min in ice. Followed by staining with specific antibodies for CD45, CD11b, Ly6G, Ly6C, F4/80, CD206, CD80 (antibody catalog number are mentioned in supplementary table S2) and fixable viability dye (65-0866-14, Invitrogen) at 4°C for 30 min. Specific isotype controls were used to determine gating. The cells were washed and fixed with 4% PFA and acquired on a ThermoFisher Attune Nxt flow cytometer. The recorded data were analyzed using FLowJO 10 software.

### Isolation of macrophages from GC muscle

The gastrocnemius muscles were minced and enzymatically digested as explained above and cells were magnetically labeled using anti-F4/80 antibody and isolated by magnetic sorting (Miltenyi Biotec).

### Quantitative RT-PCR

Total RNA was prepared from cells or tissues using Tri Reagent (Molecular Research Center Inc.). and phenol/chloroform. Reverse transcription was carried out using high-capacity cDNA reverse transcription kit (Applied biosystems) with 2µg of total RNA. The PCR was performed as per manufacturer’s protocol using ABI Prism 7000 Sequence Detection System 26 (Applied Biosystems, CA) and the QuantiFast SYBR Green PCR kit (Qiagen) for specific genes. Primer sequences for q RT-PCR are enlisted in Supplementary Table 1. The expression of genes was normalized and expressed as fold-changes relative to HPRT.

### Statistical analysis

Each experiment was repeated at least 3 times and data are presented as mean ± SEM. Comparison between two groups were analyzed by unpaired two tailed Student *t*-test. Experiments with more than 2 subgroups were analyzed by ANOVA followed by the Tukey post-hoc or Bonferroni multiple comparison analysis to specify the significance between group differences. Values of *p<0.05, **p<0.01, ***p<0.001 were considered statically significant. Statistical tests were performed using Graphpad Prism v10 (GraphPad Software, San Diego, CA).

## Author contributions

M.U.-F., T.F. and S.Y. designed the study; S.Y (major), S.V., D.A., S.N., A.D., S.K., performed/assisted research; M.U.-F., T.F., S.Y. analyzed data; V.G., discussed data and provided inputs; M.M. and S.K. performed mouse genotyping; V.G. provided materials. M.U.-F., T.F. and S.Y. wrote the manuscript.

## Acknowledgements

This work was supported by National Institute of Health (NIH) grants: R01HL160014 (to M.U.-F.), R01HL147550 (to M.U.-F., T.F), R01HL147550-S1 (to M.U.-F., T.F), R01HL133613 (to T.F., M.U.-F.), American Heart Association (AHA) Transformational project Award 22TPA971863 (to T.F.), 17POST33660754 (to D.A.); Veterans Administration (VA) Merit Review Award 2I01BX001232 (to T.F.).

---

**Figure S1:**
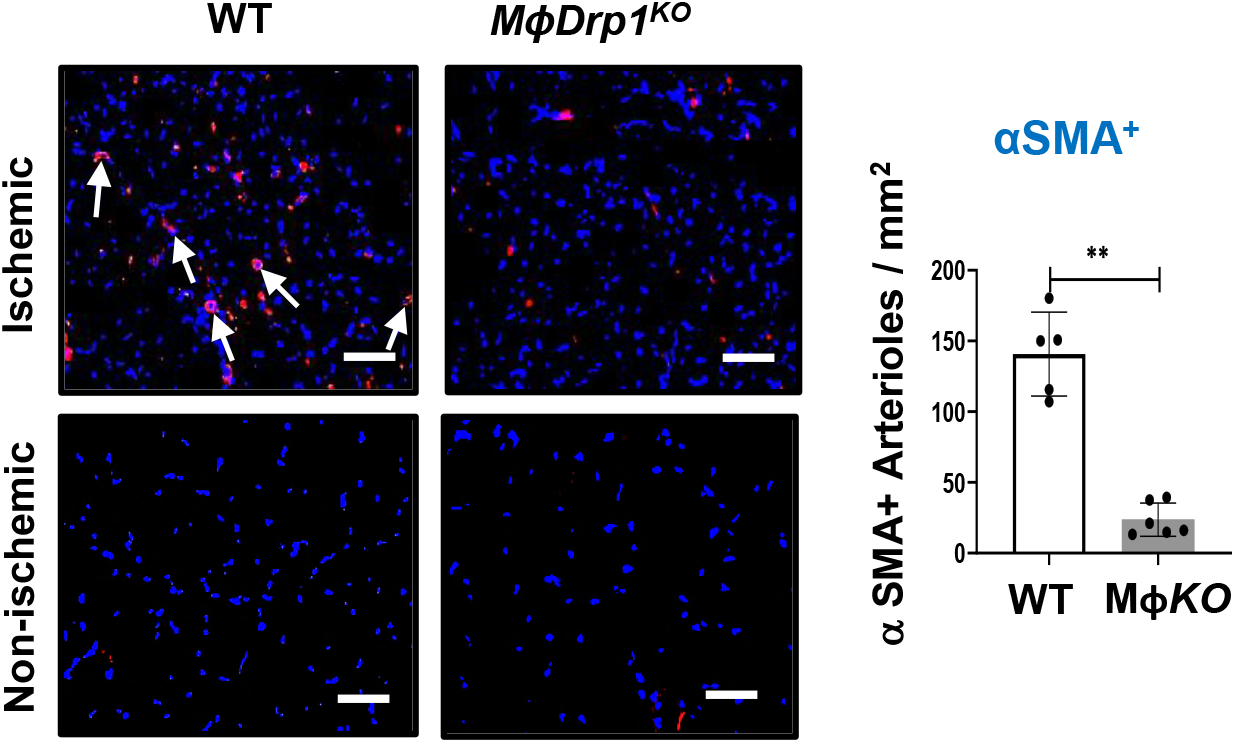
Myeloid Drp1^-/-^ mice exhibited reduced number of αSMA^+^ arterioles in ischemic muscles after HLI. Immunofluorescence images (left) and quantification (right) of α-smooth muscle cells (SMA^+^) (arterioles) in GC muscle at day 21 post-HLI. scale bar=20 µm. n=5-6. Data are mean ± SEM. **\*\***p<0.01.

**Figure S2:**
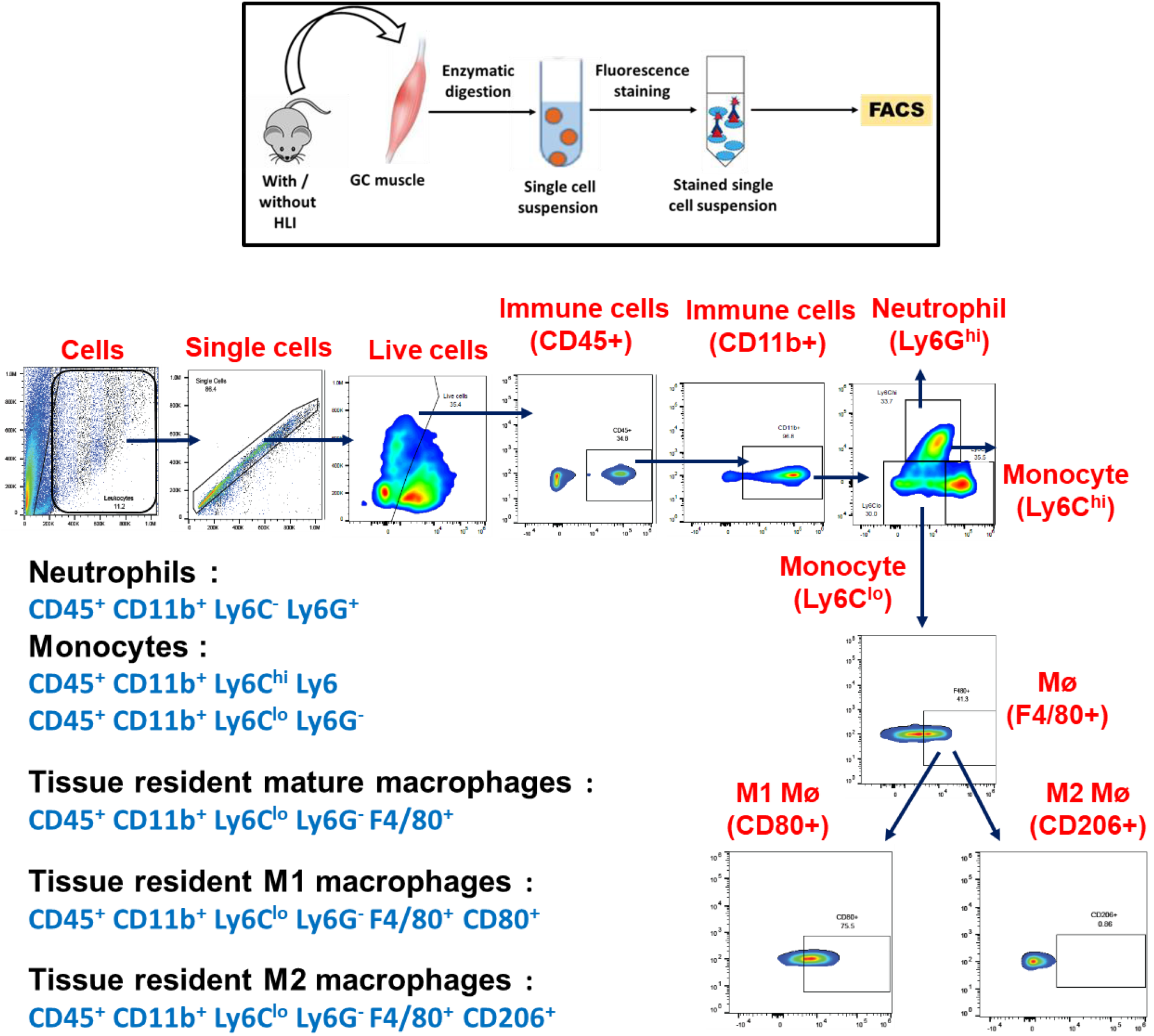
Schematic representation of gating strategy for flow cytometric based immunophenotyping of gastrocnemius muscle. Gating strategy for flow cytometry based analysis of neutrophils, monocytes and naïve (M0) macrophages, pro-inflammatory M1 and anti-inflammatory M2 macrophages isolated from ischemic GC muscle at the indicated time after HLI. FSC-A and SSC-A based cells were gated for singlets, live cells followed by CD45^+^ and CD11b^+^ double positive cells (leukocytes). Neutrophils (Ly6G^+^), monocyte (Ly6C^hi^), naïve M0 Macrophages (Ly6C^lo^ F4/80^+^), M1 macrophages (Ly6C^lo^ F4/80^+^ CD80^+^) whereas M2 macrophages (Ly6C^lo^ F4/80^+^ CD206^+^) were quantified.

**Figure S3:**
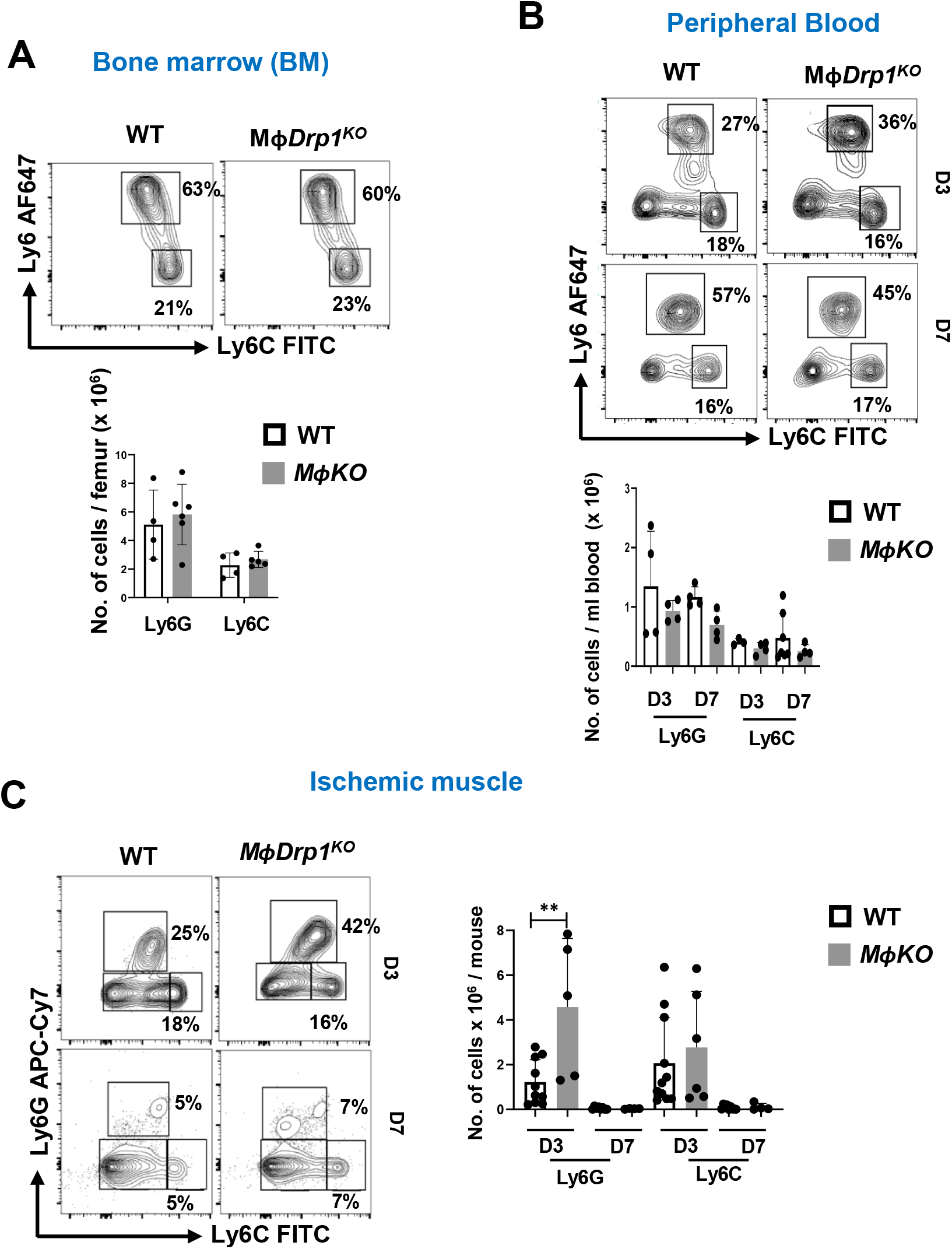
Neutrophils and monocytes in peripheral blood, bone marrow, and ischemic muscles in WT and *MɸDrp1^KO^* mice after HLI. Representative flow cytometry contour plots and quantification of numbers of Ly6G^hi^ neutrophils and Ly6C^hi^ monocytes in bone marrow (A), peripheral blood (B) at day 3 post-HLI as well as in ischemic GC muscle at days 3 and 7 post-HLI in WT and *MɸDrp1^KO^* mice. Data are mean ± SEM. n=3-5. **p<0.01.

**Figure S4:**
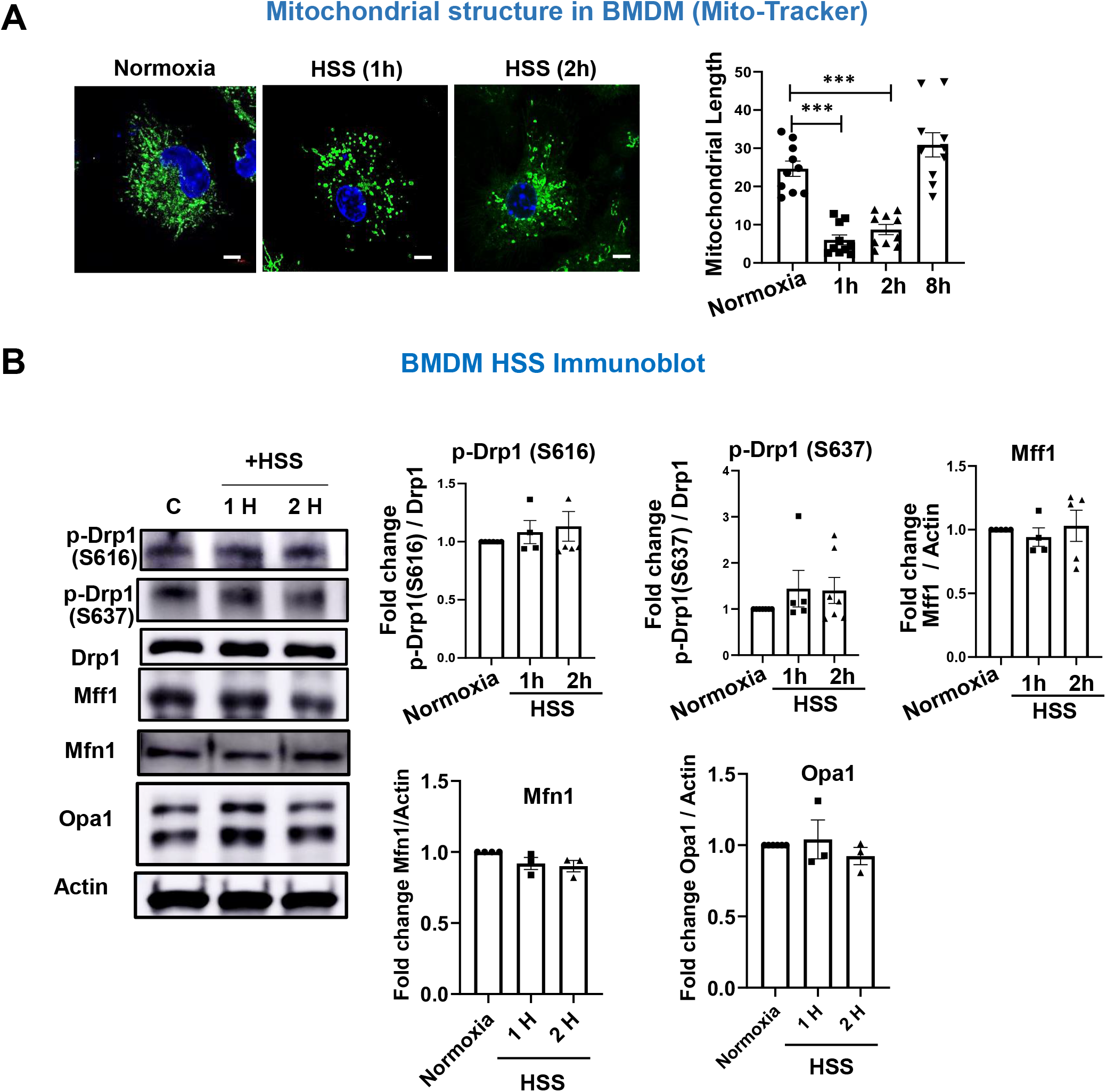
BMDM exposed to HSS induced mitochondrial fission without altering expression of key mitochondrial fission/fusion protein. **A.** Analysis of mitochondrial dynamics in WT BMDM stained with Mito-tracker green exposed to HSS for indicated time (left). Right panel shows quantification of mitochondrial length measured by image J (right). Scale bar=5μm. ***p<0.001. B. Western blot analysis for p-Drp1-S616, p-Drp1-S637, total Drp1, Mff1, Mitofusin 1 (Mfn1), OPA1 and actin (loading control) in WT-BMDM exposed to HSS for indicated time. Right panels are quantification. Data are mean ± SEM. n=3-5.

**Figure S5:**
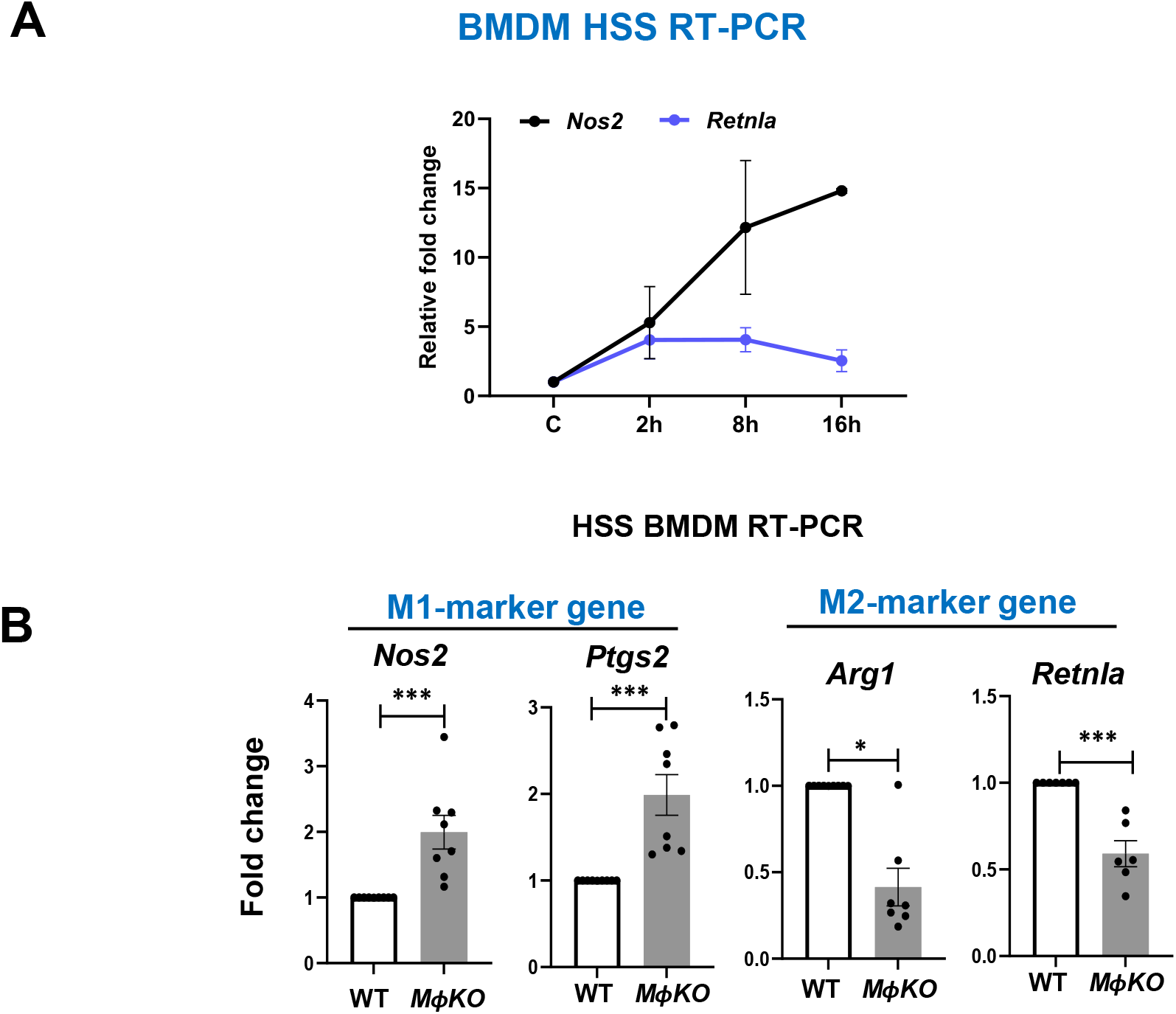
Drp1^-/-^ BMDM under HSS exhibited increase in M1-marker genes and decrease in M2-marker genes. A. Quantitative RT-PCR (relative to *Hprt*) of M1-gene *Nos2* and M2-gene *Retnla* mRNA levels in WT-BMDM exposed to HSS at indicated time. Data are expressed as fold increase vs. normoxic control. B. M1 markers (*Nos2* and *Ptgs2)* and M2 markers (*Arg1* and *Retnla)* mRNA levels in WT and *Drp1^KO^* BMDM under HSS for 8 h. Data are mean ± SEM. n=3-7. *p<0.05, ***p<0.001.

**Figure S6:**
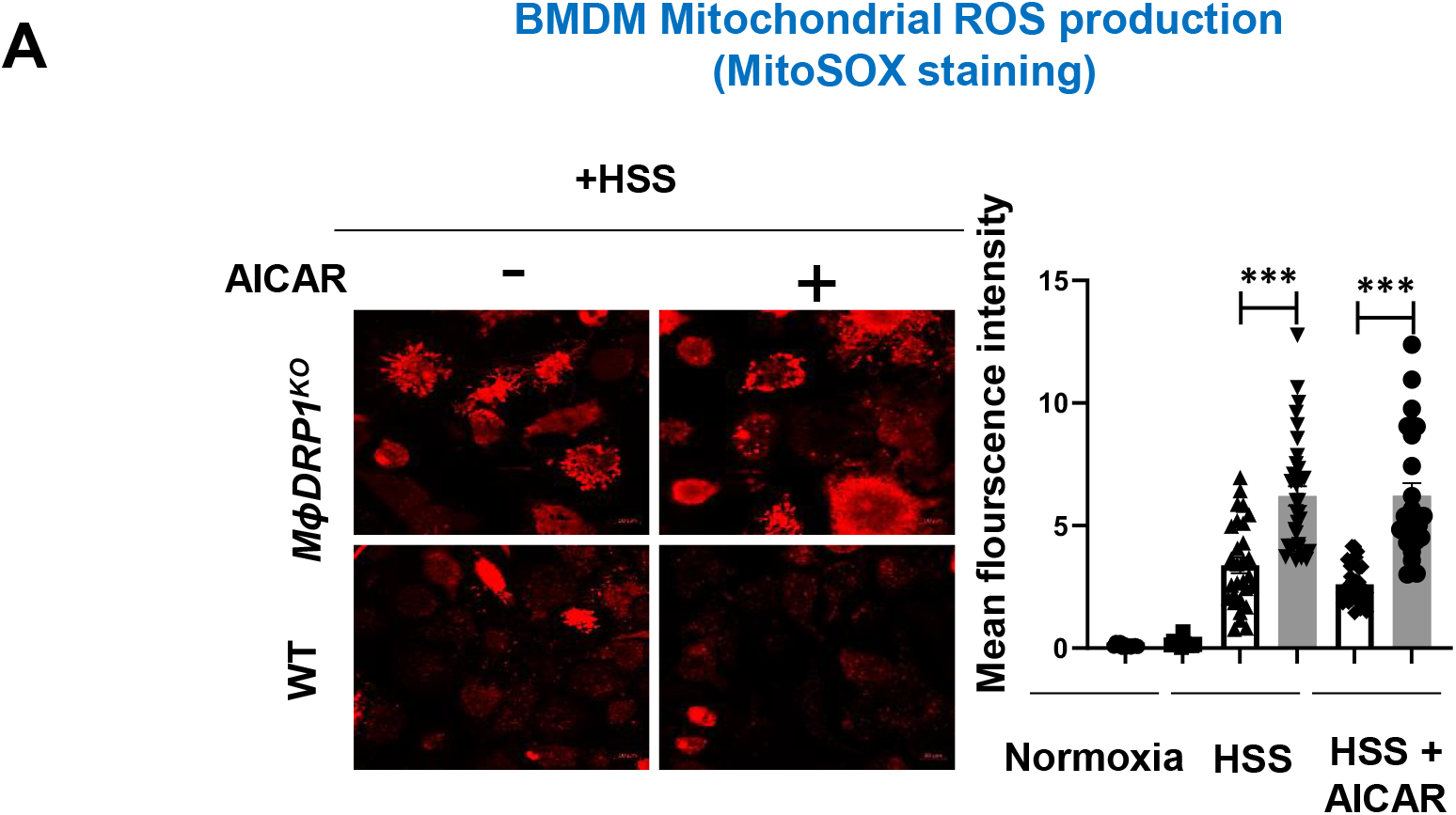
Excess mitoROS production in Drp1^-/-^ BMDM under HSS is not suppressed by AMPK activator AICAR. Left, Effects of AMPK activator AICAR (pre-treatment for 2h at 100 μM) on mitochondrial ROS measured by MitoSOX in WT and Drp1^-/-^ BMDM under HSS for 2 h. Right panel shows the quantification of mean fluorescence intensity using ImageJ. Scale bars=5μm. ***p<0.001.

**Table S1:**
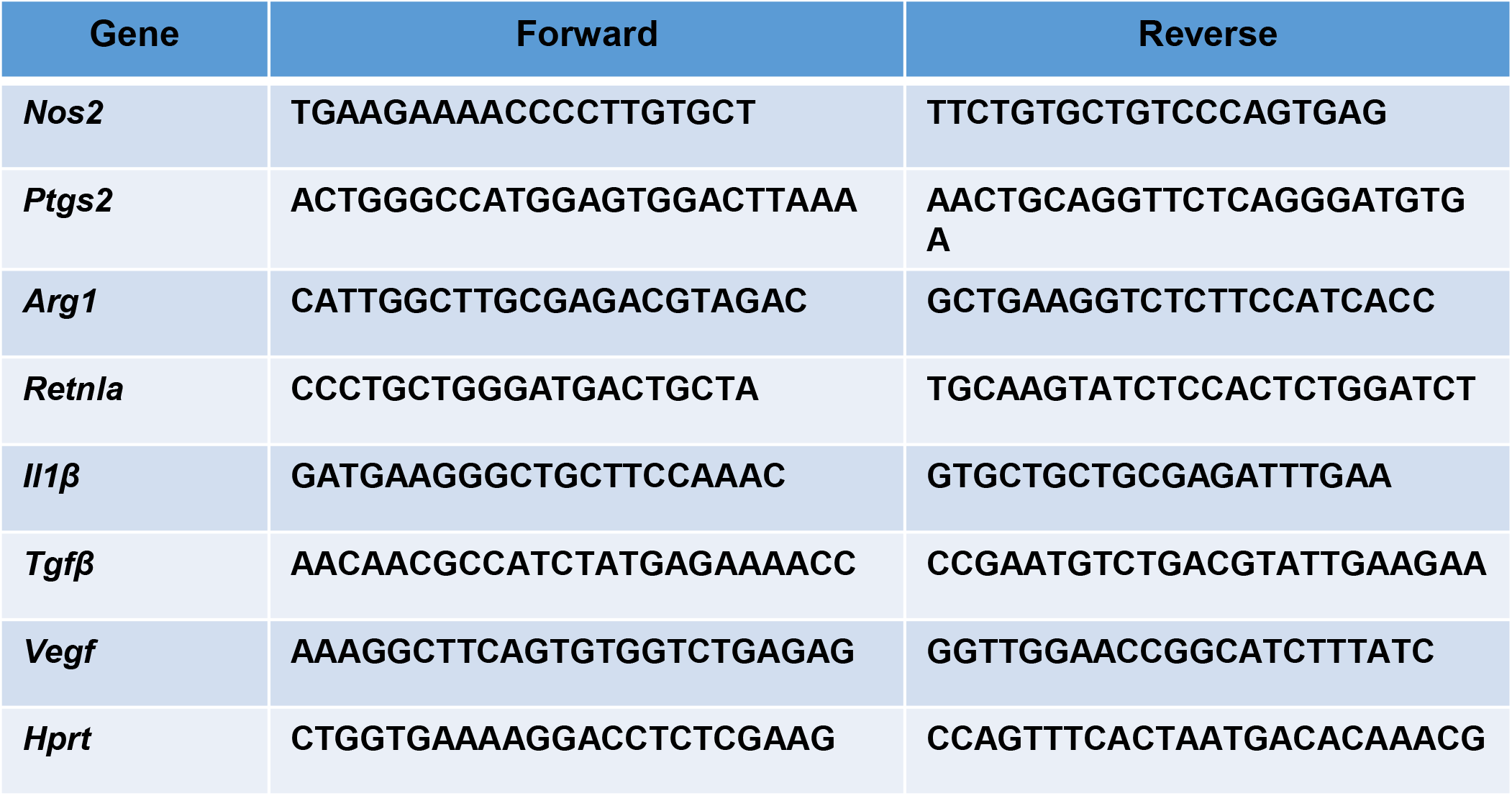
Sequences of primers used for RT-PCR.

**Table S2:** Antibodies used for flow cytometry based immunophenotyping.

| Antibody | Source | Catalog No. |
| --- | --- | --- |
| PE anti mouse CD45 | Biolegend | 147712 |
| PE/Cyanine anti mouse CD11b | Biolegend | 101216 |
| PE anti mouse CD11b | Invitrogen | 12-0112-82 |
| FITC anti-mouse Ly-6C | BD Bioscience | 553104 |
| Alexa 647 anti mouse Ly-6G | Biolegend | 127610 |
| PE/Cyanine anti mouse Ly-6G | Biolegend | 127624 |
| PerCP-Cyanine5.5 anti mouse F4/80 | Invitrogen | 45-4801-80 |
| APC anti mouse F4/80 | Biolegend | 123116 |
| BV421 anti mouse CD80 | BD Bioscience | 562611 |
| BV605 anti mouse CD206 | Biolgend | 141721 |

